# Refined movement analysis in the Staircase test reveals differential motor deficits in mouse models of stroke

**DOI:** 10.1101/2023.10.23.563529

**Authors:** Matej Skrobot, Rafael De Sa, Josefine Walter, Arend Vogt, Raik Paulat, Janet Lips, Larissa Mosch, Susanne Mueller, Sina Dominiak, Robert Sachdev, Philipp Böhm-Sturm, Ulrich Dirnagl, Matthias Endres, Christoph Harms, Nikolaus Wenger

**Affiliations:** Department of Neurology with Experimental Neurology, Charité – Universitätsmedizin Berlin, Charitéplatz 1, 10117 Berlin, Germany; QUEST Center for Transforming Biomedical Research, Berlin Institute of Health (BIH), Berlin, Germany; Center for Stroke Research Berlin, Charité - Universitätsmedizin Berlin, Germany; NeuroCure Cluster of Excellence and Charité Core Facility 7T Experimental MRIs, Charité-Universitätsmedizin Berlin, Berlin, Germany; Institute of Biology, Humboldt University of Berlin, Berlin, Germany; Sussex Neuroscience, School of Life Sciences, University of Sussex, Brighton BN1 9QG, UK; DZHK (German Center for Cardiovascular Research), Berlin Site, Germany; DZNE (German Center for Neurodegenerative Disease), Berlin Site, Germany; DZPG (German Center of Mental Health), Berlin Site, Germany

**Keywords:** machine learning, motor deficits, rodent models, stroke, translational research

## Abstract

Accurate assessment of post-stroke deficits is vital in translational research. Recent advances in machine learning provide unprecedented precision in quantifying rodent motor behavior post-stroke. However, the extent to which these tools can detect lesion-specific upper extremity deficits remains unclear. Using proximal middle cerebral artery occlusion (MCAO) and cortical photothrombosis (PT), we assessed post-stroke impairments in mice through the Staircase test. Lesion locations were identified using 7T-MRI. Machine learning was applied to reconstruct kinematic trajectories using *MouseReach*, a data-processing toolbox. This yielded 30 refined outcome parameters effectively capturing motor deficits. Lesion reconstructions located ischemic centers in the striatum (MCAO) and sensorimotor cortex (PT). Pellet retrieval was altered in both cases but did not correlate with stroke volume or ischemia extent. Instead, cortical ischemia was characterized by increased hand slips and modified reaching success. Striatal ischemia led to progressively prolonged reach durations, mirroring delayed symptom onset in basal ganglia strokes. In summary, refined machine learning-based movement analysis revealed specific deficits in mice after cortical and striatal ischemia. These findings emphasize the importance of thorough behavioral profiling in preclinical stroke research to increase translational validity of behavioral assessments.

## Introduction

Worldwide, stroke is one of the leading causes of long-term disability.^1^ About three out of four stroke survivors suffer from acute upper limb deficits.^2^ A central objective of translational stroke research is to better understand the prognosis of motor recovery and find personalized treatment strategies.

One important predictor of post-stroke recovery is the severity of the initial deficit.^3–5^ After severe paresis, only half of stroke survivors regain meaningful levels of upper limb function.^6^ The recovery of arm movements benefits to some extent from specialized forms of physiotherapy, such as constraint-induced movement therapy, mental practice, robotics, or EMG biofeedback.^7^ Yet, deficits in fine-skilled hand movements persist across treatment modalities.^7^

Lesion location is a second important factor predicting post-stroke recovery. In humans, isolated cortical stroke displays a higher degree of upper limb recovery compared to subcortical stroke in the corona radiata, basal ganglia, or thalamus.^8^ Different lesion locations are characterized by heterogeneous symptoms in humans. For example, a stroke along the corticospinal tract leads to sudden-onset hemiparesis.^9,10^ In contrast, symptoms after isolated basal ganglia infarct can manifest with prolonged aggravation, for weeks and months, with movement disorders such as dystonia or hyperkinesia.^11^ In rodent stroke models, it is not known whether distinct behavioral deficits will similarly relate to lesion location.

Typical assessments of forelimb function in stroke models capture global behavioral parameters such as pellet retrievals, percentage of limb use in a cylinder, or paw placement on a ladder.^12,13^ These global scores measure changes in performance but lack the capacity to discriminate cognition from sensorimotor function or sickness behavior. When using such global metrics, it is further difficult to distinguish true recovery from the development of compensatory movement strategies after stroke.^14–16^ These limitations have been readily recognized by interdisciplinary expert consortia (STAIR, SRRR), and it has been recommended to integrate kinematic analysis as broadly as possible for the assessment of sensorimotor function in preclinical stroke research.^17,18^

Machine learning has opened compelling avenues for studying complex motor behaviors in rodents without the need for marker-based motion tracking.^19–26^ These tools have helped unravel neural circuits responsible for coordinated hand function in physiological states in mice.^27,28^ More recently, these tools have been successfully adopted for the analysis of gait in mice following cortical photothrombosis.^29^ Given these new technological opportunities, deep behavioral profiling^30^ is about to reveal its full potential for the improvement of translational stroke research.

Here, we examined whether machine learning can refine the analysis of skilled forelimb use and potentially distinguish lesion-specific deficits in mice after stroke. We chose to work with the Staircase test, a well-defined environment for studying coordinated movement.^18,31^ In the task, animals enter into a small reaching chamber to then dynamically grasp food pellets that are placed on a staircase at different distances. During reaching attempts, animals maintain their body stability through an isometric grip with their opposite hand on an elevated platform. Instead of relying on traditional pellet counts, we reconstructed bilateral hand and food pellet trajectories during task execution with the software package *DeepLabCut.*^19^ Next, we developed *MouseReach*, an automated data-processing toolbox, to derive meaningful parameters for the quantification of post-stroke motor performance. *MouseReach* algorithms automatically detected reaching attempts towards the pellets and vertical slips of the hand from the stabilizing platform. Reaching attempts were further classified into reaches without pellet contact, successful pellet removals or pellet drops during retrieval. The toolbox was further used to analyze kinematic features of reaching attempts, yielding a total parameter set of 30 outcome parameters for refined motor deficit quantification. Application of *MouseReach* accurately captured the time course of differential symptom manifestation following a middle-cerebral artery occlusion (MCAO) or cortical photothrombosis (PT) in mice.

## Methods

### Experimental design

We used 8–10-week-old C57BL/6N (20–26 g) mice from Charles River Laboratories in our experiments. Animals were housed in enriched home cages in groups of three with a regular day-night cycle. A total of 12 mice (6 females and 6 males) were included in the MCAO group and 10 mice (5 females and 5 males) in the PT group. A subcutaneous chip was implanted to allow for daily measurements of animal identity and body temperature. One female mouse was later excluded from the PT group due to anesthesia-related death, and two mice were excluded from the MCAO group after not learning the Staircase task prior to the experimental stroke. Three days before the start of the Staircase training, animals received restricted access to food. Food access was provided ad libitum for 3 hours a day, immediately after the Staircase exposure. For the rest of the day-night cycle, an additional 1.1 g of food per animal was added to each home cage. Daily training and weekly functional assessments were performed between 8 and 11 a.m. In the case of weight loss greater than 5% of bodyweight compared to baseline, the amount of food was increased. Food restriction was suspended for weight loss greater than 10%. Water was available ad libitum. Experiments were conducted after approval by the Berlin State Office for Health and Social Affairs (LAGeSo) under the licenses G0108/20 and G0343/17.

### Staircase training

Following two weeks of handling, the animals started training in the Staircase for 30 minutes per day to learn how to grab and eat dustless sugar pellets (20 mg sucrose, TestDiet) with the left and right hands. After one week, the duration of the reaching session was reduced to 20 minutes. The remaining pellets in each Staircase box were counted after every trial and reported as the ‘traditional pellet count’. Mice that removed less than two of eight available pellets per side were excluded from the experiment prior to the induction of ischemia. Animals participated daily in the Staircase test, except for the day of stroke intervention.

### Stroke models

For all surgeries, mice were kept under 1%–2% isoflurane anesthesia. Analgesia was achieved with an s.c. injection of carprosol (5 mg/kg). Body temperature was maintained at 37°C on a heating pad. For photothrombosis (PT), mice were transferred to a stereotactic frame (Kopf Instruments) to receive a unilateral ischemic lesion of the left sensorimotor cortex, as previously described.^32^ In brief, the skull was exposed by a midline incision of the scalp. An opaque, reflective template with a defined opening (3 mm wide, 5 mm long) was aligned to the midline over the left sensorimotor forelimb and hindlimb areas: −2 to +3 mm A/P, 0 to −3 mm M/L related to bregma. Five minutes after an intraperitoneal injection of 200 µL of Rose Bengal (10 mg/ml in 0.9% NaCl; Sigma-Aldrich), the skull was illuminated with a light source (Zeiss, CL 1500 HAL, 150W, 3000K) that was securely placed on top of the skull for 15 minutes. For MCAO, we used a transient, 45-minute occlusion of the left middle cerebral artery.^33^ Ischemia was induced by introducing a 7–0 silicon rubber-coated MCAO suture with a coating length of 9–10 mm (monofilament 7019910PK5Re, Doccol Corp., Sharon MA, United States) into the left internal carotid artery and advancing it up to the anterior cerebral artery, thereby occluding the middle cerebral artery (MCA). The filament was later withdrawn after an occlusion time of 45 min. After both procedures, the animals were allowed to recover in a heated cage for a minimum of 1 hour. Following surgery, all animals had access to wet food. If any signs of dehydration occurred, animals were treated with additional subcutaneous saline injections.

### 7T-MRI

T2-weighted images were acquired one day after stroke in the MCAO and PT groups. An experienced researcher used ANALYZE software (v5.0, AnalyzeDirect, Overland Park, KS, USA) to segment the hyperintense lesion. MR-images and the lesion mask were registered on the mouse Allen Brain Atlas using the in-house developed MATLAB toolbox ANTx2 (https://github.com/ChariteExpMri/antx2). Incidence maps expressing the percentage of animals with a lesion in a voxel were plotted for each group in atlas space, and edema-corrected lesion volumes were calculated, as previously described.^34^ Brain region-dependent infarct size was defined as the percentage of lesioned isocortex or striatum.

### Multi-Staircase setup

To reduce overall experimental workload for testing multiple animals, we designed a Staircase setup for simultaneous video recordings in four mice (Figs. 1 and S1). For this, four Staircase boxes were positioned in a 2×2 grid using a customized positioning platform. The platform was designed to guide the accurate placement of each Staircase box, and to minimize time for camera calibrations across days. Each Staircase box consisted of two sub-compartments, a resting chamber, and a reaching chamber containing a dual 8 well staircase (Fig. S1A). We placed one sugar pellet per staircase well to help software algorithms determine when wells were either full or empty. At the entry to the reaching-chamber, each Staircase box was equipped with an infrared break-beam sensor (Adafruit) to detect periods of reaching activity and to prevent overly large video file sizes. Beam break crossings triggered the recording of two high-speed cameras (ACE GigE, Basler, Germany) that were positioned on either side of the Staircase setup. Videos were recorded at a time resolution of 100 frames per second (fps) with a spatial resolution of 640×480 pixel. The camera image was centered on four opposing reaching chambers for parallel recordings of multiple mice (Fig. 1A). In real-time, a customized trigger box (Fig. S1B) integrated information from all beam breaks and performed a logical AND operation to avoid repeated start signals when several animals entered the reaching chambers. Individual recordings stopped when all mice returned to their resting chambers.

**Figure 1.**
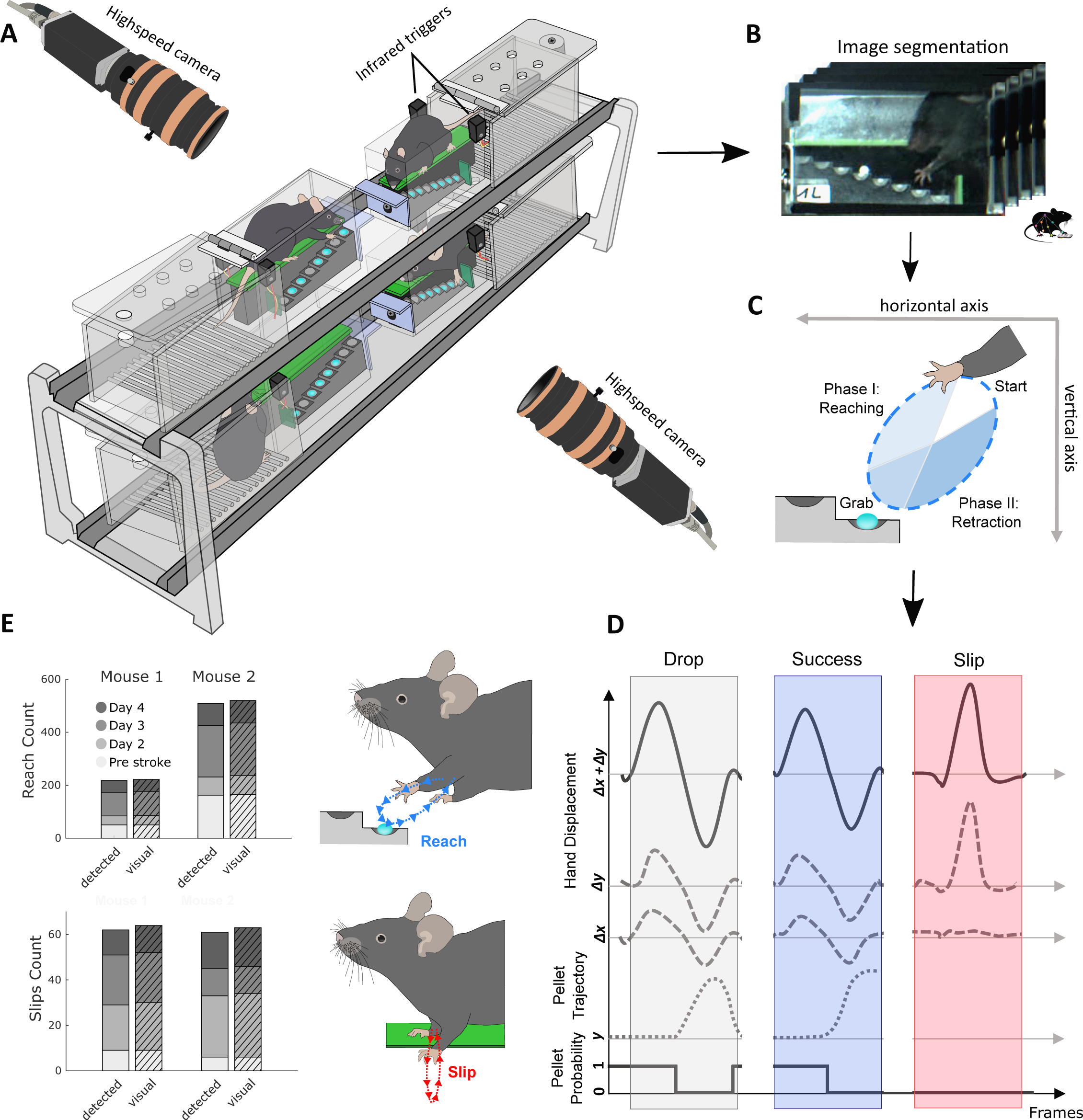
Setup for refined movement analysis and validation of algorithm performance. (A) Four staircase boxes are simultaneously recorded using two highspeed cameras that are triggered by infrared beams, when mice enter the reaching chambers. (B) Images from individual video-frames are segmented and streamlined for machine learning based motion tracking of hand and pellet trajectories in individual Staircase boxes. (C) The direction of hand movement during reach cycles defines movement towards (reach-phase) and away from a target sugar pellet (retraction-phase). (D) Successful and unsuccessful reaches are detected based on logical threshold operations for hand and pellet displacements. Slips were characterized by sudden vertical movements, independent of pellet information. Probability denotes the probability of pellet existence in a staircase well. Δx and Δy: horizontal and vertical hand displacement; y: vertical pellet position. (E) The validation of automatically detected reaches and slips shows high degrees of accuracy in comparison to visual annotation by blinded raters. Data in E present results for two mice from the MCAO and PT groups, before and after stroke on days 7, 14, and 21.

### MouseReach Software

The toolbox *MouseReach* consists of a pipeline for data-file processing, event classification, and kinematic analysis that generates a set of 30 outcome parameters for each side of the body (Table S1). The toolbox source code is freely available for download from online repositories under the link: https://github.com/Wenger-Lab/MouseReach. Algorithms are both applicable to videos from single and multi-Staircase setups.

#### Data-file processing

Prior to machine learning, we segmented individual videos into four subsections, each showing the image of one of the four reaching chambers (final resolution of 320×240 pixel per subsection). Next, we matched video subsections to blinded animal identities from manual entries. The resulting dataset was used to train a neural network with the software package DeepLabCut.^19^ A total of approximately 500 video frames were manually marked as a training set. Video stacks were then automatically streamed to DLC for marker less tracking of hand and pellet trajectories in 2D. A custom true-or-false filter discarded non-physiological trajectory jumps that remained present in the DLC neural network despite optimized training. Non-physiological trajectory jumps were defined as deviations of six times the standard deviation for the 2D trajectories. Missing data points that typically affected single video frames were filled with the x and y mean values of the two nearest, correct video frames. All trajectories were then smoothened with a moving mean of 10 frames to reduce trajectory noise.

#### Event Classification

Both reaching attempts and hand slips were identified based on hand velocity when a successive pair of maximal and minimal velocities exceeded an empirically determined threshold (1.5 pixels per frame). The classification of events into either reaching attempts or slips was achieved by calculating the ratio of hand displacement in horizontal (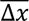) and vertical (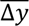) directions. A ratio smaller than 1.5 identified slips. Detection of pellet removals was achieved by computing a frame-by-frame binary signal based on the probability of pellet existence in each of the eight staircase wells. A pellet removal was detected when the probability dropped below 99%. The pellet removal was then matched in time with the corresponding reaching attempt. Starting from the frame of pellet removal, a common trajectory of the pellet and the hand was monitored. If the pellet disappeared prior to the next reaching attempt and no pellet drop was detected, the reaching attempt was counted as successful (reaches_successful_). Reaching attempts were counted as unsuccessful if a pellet drop was detected during or after a reach. The target for each reaching attempt was calculated by finding the closest staircase well to the inversion point of hand movement in 2D. Using this information, we defined a new coefficient of reaching success (k_success_):

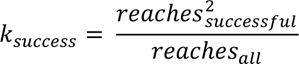

The term *reaches_all_* represents the total count of reaches towards staircase wells, that were not yet emptied by the mouse. *reaches_successful_* counts all successful pellet retrievals from the set of *reaches_all_.* In the formula, we square the term *reaches_successful_* to reward mice with a higher score when they retrieve a higher number of pellets. Together, the event classification algorithms resulted in five outcome parameters of motor performance: success coefficient and number of success events, pellet events, reach events and slip events.

### Kinematic Analysis

During analysis, pixels and frames were converted to centimeters and seconds, using staircase dimensions from the video and frame rate. Reaching movements were further divided into two discrete phases: a ‘reach’ phase towards or a ‘retraction’ phase away from the pellet (Fig. 1c). The start, midpoint, and end of a single reaching attempt were calculated based on zero crossings of hand velocity. For each of the three categories (full reaching attempt, reach, and retraction phases), we calculated distance, duration, average velocity, and average acceleration. For instantaneous velocity and acceleration, we also determined the respective minima and maxima. For slip events, we calculated ‘slip depth’ as the distance of the vertical hand displacement from the green staircase platform (Fig. 1E). All calculations were performed as Euclidian metrics. For hand displacement during reaching (***Δx+Δy***), the total path length ***s*** was calculated as

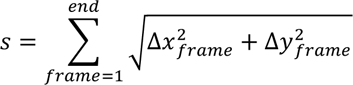

*with frame = 1 and end* being the starting and end frame of each detected reaching event. Frame-by-frame speed ***v*** and acceleration ***a*** were quantified as time derivatives:

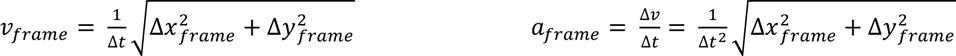

with Δt being the time interval per video-frame (10 ms at a framerate of 100 Hz).

Together, these calculations resulted in a total of 30 outcome parameters for event quantification and kinematic analysis (Table S1).

### Statistics

The planning of group sizes was performed prior to experiments, based on expected effect sizes for stroke induced changes in pellet count, calculated separately for each stroke intervention (for MCAO: effect size 0.85, alpha error 0.05, resulting group size = 11. For PT: effect size: 0.95, alpha error: 0.05, resulting group size = 9). Experimenters and raters were blind to group allocation. All presented analysis was performed by automated algorithms. To find outcome parameters that maximize the difference between stroke models and timepoints of observation, we used linear discriminant analysis (LDA). Normal distribution was tested with the Kolmogorov-Smirnov test. For statistical quantification of individual outcome parameters, we performed a nonparametric, two-way repeated measures ANOVA for the factors stroke model and timepoint.^35^ For post-hoc analysis, pairwise comparisons were performed using the Wilcoxon rank sum test (for comparisons between different stroke groups), and the Wilcoxon signed rank test (for paired comparisons within the same stroke group at different timepoints). Due to the novelty of calculated functional parameters, statistical comparisons are of an exploratory nature, and p-values have not been adjusted for multiple testing. In the figures, all bar graphs are reported using mean ± standard deviation (SD) and box-and-whisker plots are displayed using the Tukey method. Correlations between lesion volume and symptoms were calculated with linear regressions and Pearson correlation coefficients. The accuracy, sensitivity, and specificity of automated event classifications were calculated in comparison to manual annotations as a ground truth. The manual annotation was performed with a custom-programmed graphical user interface in MATLAB by blinded raters.

## Results

### Diagnostic performance of *MouseReach* for movement classification

We first validated the performance of our automated movement classifications in comparison to manual annotations. Automated classification was based on the information of two high-speed video cameras (framerate 100Hz) that recorded mouse behavior from either side of the multi-Staircase setup (Fig. 1A). Images of each video frame were then automatically segmented into four individual chambers and streamlined to the machine learning software, *DeepLabCut*, for tracking of hand and pellet trajectories (Fig. 1B). Hand trajectories were used to detect reaching attempts and vertical hand slips. Reaching attempts were further subdivided into a reaching phase towards and a retraction phase away from the pellet (Fig. 1C). Targeted pellets were identified by the nearest pellet to hand at the inversion of the two reaching phases. Reaching attempts were further classified into reaches with no pellet contact, successful pellet retrieval or unsuccessful retrieval when the target pellet dropped onto the staircase floor prior to the next reaching event. Sudden downward shifts of the hand, limited to the vertical axis, were detected as hand slips from the staircase platform (Fig. 1D).

After establishing processing algorithms, we validated the performance of automated event classification for four event categories: reaching attempts, hand slips, pellet removals and final pellet count in the box (Figs. 1E and S2). As a ground truth, one mouse from each stroke group was randomly picked for manual quantification of all events on four recording days, including one day of pre-stroke behavior. Reaching attempts were correctly identified with an accuracy of 98.3%, a sensitivity of 98%, and a specificity of 100%. Slip detection showed a performance of 99.5%, 96.9%, and 100%, respectively. Pellet removals were identified at 99.8%, 98.5%, and 99.9%. This high degree of accuracy confirmed the suitability of the automated algorithms for the analysis of stroke deficits at group levels. The high number of recorded events (i.e., up to 200 reaching attempts per animal per day) rendered manual annotation very laborious (approx. 4 hours of annotation per animal per recording day), further highlighting the importance of automated processing.

### Quantification of lesion volume and location with MRI morphology

We next investigated the distribution of lesion locations in the 45-min MCAO and PT stroke models, using MRI-based morphological reconstructions. Before the stroke intervention, all animals were subject to 4 weeks of food restriction and 3 weeks of daily Staircase training (Fig. 2A). Video recordings in the Staircase test were taken on day 4 before and days 7, 14, 21 after stroke. MRI measurements were performed on day one after the stroke. Stroke incidence maps were reconstructed according to the Allen Brain Atlas (Fig. 2B). MCAO and PT resulted in an average lesion volume of 22.99 mm^3^ (±13,2 SD) and 34.03 mm^3^ (±9,8 SD), respectively, with no significant difference for group comparison (Fig. 2C). Lesions were primarily present in the cortex and striatum for both stroke groups, with only minimal lesion volume in other brain areas. MCAO predominantly induced lesions in striatum, with partial cortical involvement (37.75% striatal vs. 16.2% cortical, p < 0.01). In contrast, PT generated small striatal and large cortical lesions (4.97% vs. 46.19%, p < 0.001). Lesion volumes in the remaining brain areas accounted for 2.94% in PT and 3.28% in MCAO. Due to their structural predominance, we used striatal and cortical lesions for further correlation analysis in this study.

**Figure 2.**
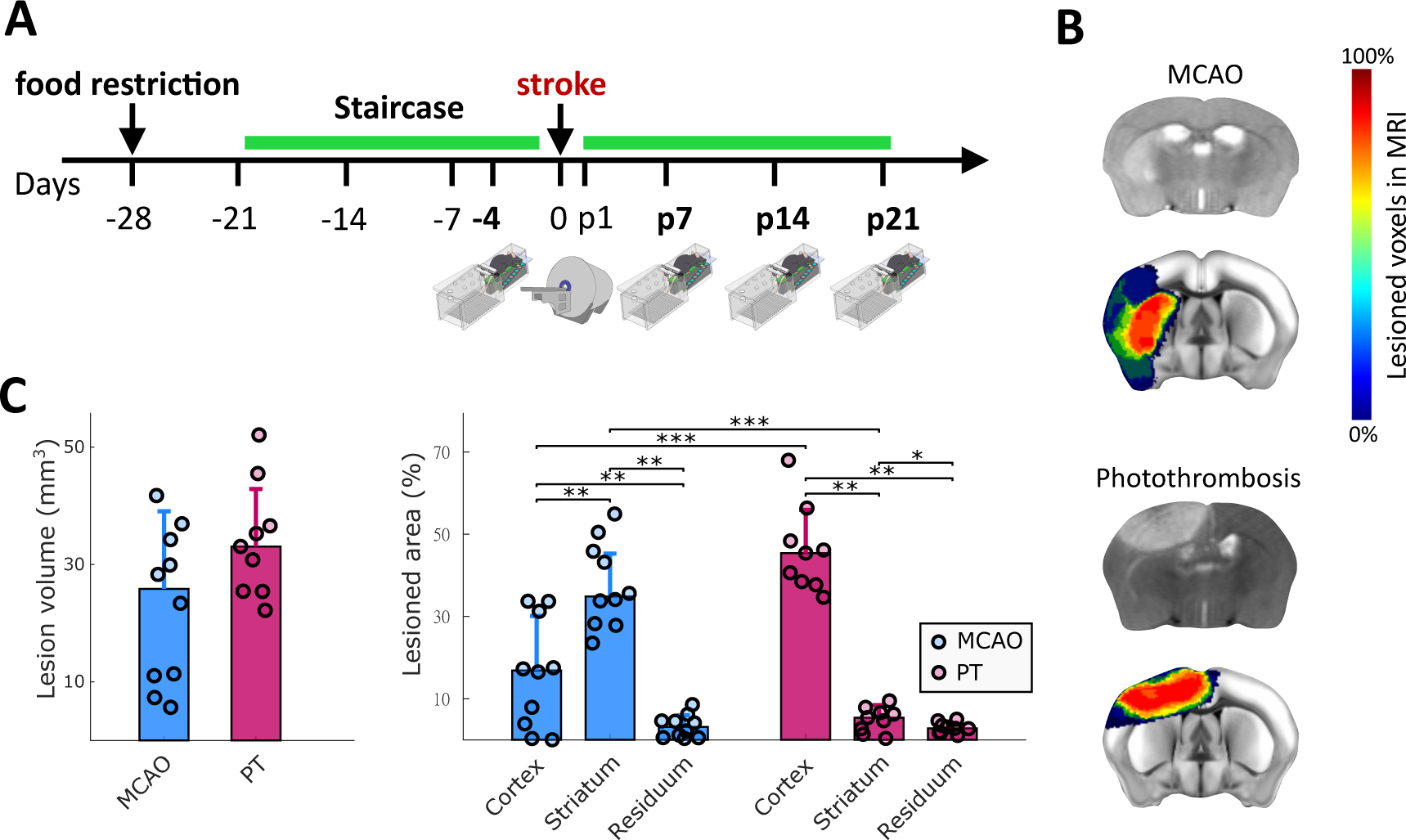
Experimental timeline and quantification of lesion volumes and locations. (A) Animals were trained in the staircase daily (green bar), except for the day of stroke surgery. Video recordings were performed on day 4 before, and days 7, 14, and 21 after stroke. Days before and after stroke are indicated with symbol ‘-‘ and letter ‘p’. MRI was performed on day 1 after stroke. (B) Representative MRI cross sections after MCAO and PT (grey scale images), and colored stroke incidence maps at the group level for MCAO (n = 10) and PT (n = 9) mice. Color bar indicates percentage of mice that showed lesions in individual MRI voxels. (C) Total lesion volume and percentage of lesion affecting cortical, striatal, and residual brain areas. Percentages in C are referenced to the stroke affected hemisphere. Bar graphs are reported as mean ± SD, *p<0.05, **p < 0.01, ***p<0.001.

### Stroke model specific deficits in the right hand following left sided ischemia

We next sought to identify the most relevant functional deficits following either MCAO or PT in mice. For this, we computed a set of 30 parameters for motor deficit quantification, including global parameters such as success ratio of reaching attempts, hand slip count, or kinematic features of reaching attempts (List in Table S1). To identify the most relevant features for deficit quantification, we performed a linear discriminant analysis (LDA) for supervised dimensionality reduction based on group information from stroke lesions and recording days (Fig. 3 and Fig. S4). The first linear discriminant (LD1) explained 41.5% of the variance in the dataset, and LD2 explained 24.5% (Fig 3A). Both stroke groups showed significant changes in LD1 and LD2 scores (p < 0.001). The most prominent differences in LD1 were observed for MCAO on day 21 (p < 0.01) and PT on day 7 (p < 0.01) when compared to pre-stroke values (Fig. 3B and S3). Similar effects were observed for LD2 (Fig. S4A). Factor loadings on LD1 revealed that stroke-induced behavioral changes were predominantly related to global motor deficits, movement speed of reach kinematics but not absolute kinematic distances (Fig. 3C). Together, this multifactorial analysis revealed a predominance of early deficits after PT vs. accumulating late deficits after MCAO.

**Figure 3.**
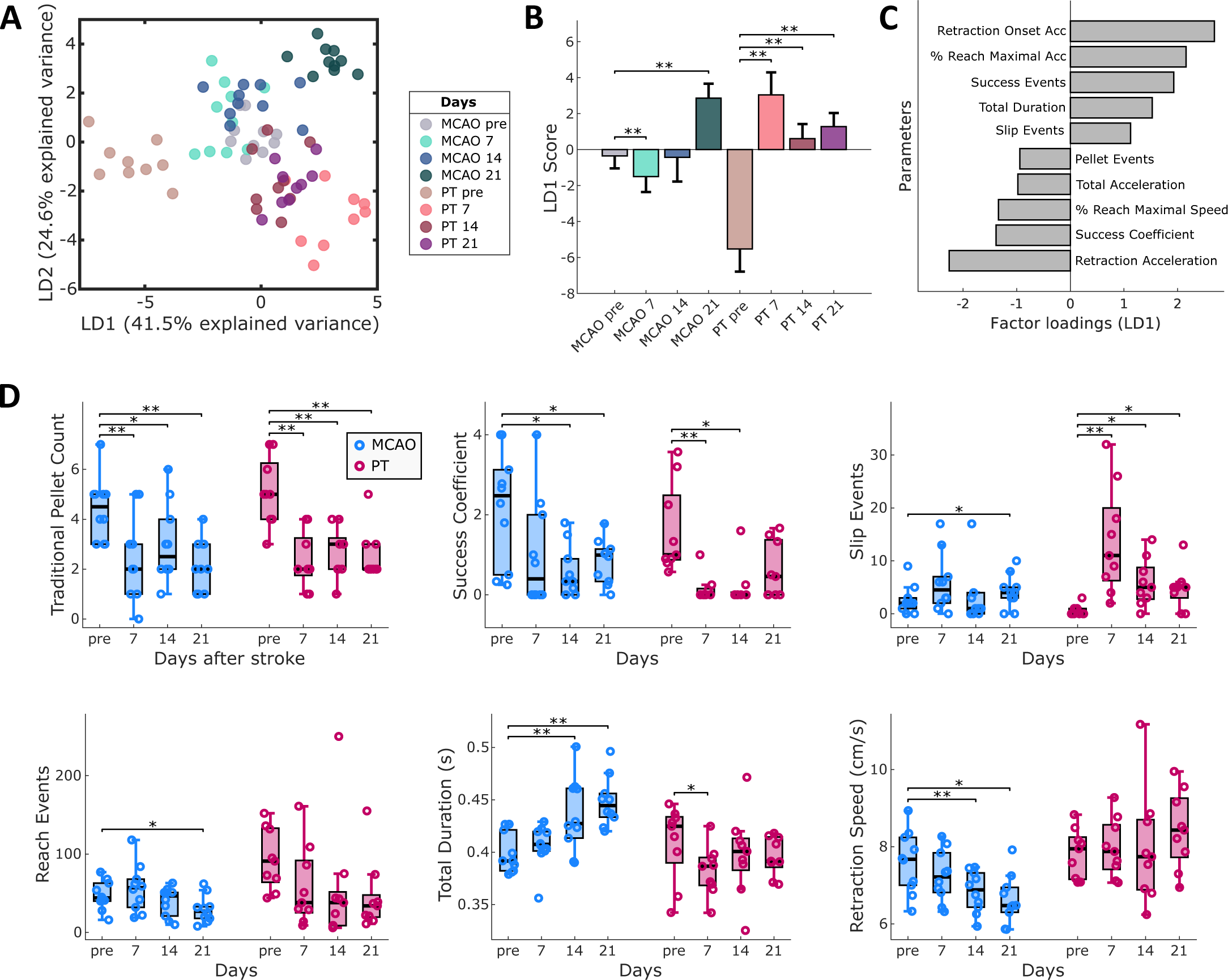
Lesion-specific deficits in the contra-lesional hand following MCAO or PT in mice. (A) Linear discriminant analysis (LDA) performed on 30 outcome parameters separates stroke groups and recording days. (B) LD1 scores show significant changes in post-stroke behavior for MCAO and PT groups. (C) The top ten factor loadings on LD1 reveal contribution of speed-related and global parameters to post-stroke deficits. (D) Comparison of post-stroke deficits using traditional quantification (pellet count) vs. refined outcome parameters. Bar graphs are reported as mean ± SD and box plots with Tukey method, *p<0.05, **p < 0.01.

At the level of individual outcome parameters, traditional pellet count successfully captured post-stroke deficits in both stroke groups, including days 7, 14, and 21 (Fig. 3D). In contrast, the success coefficient showed significant deficits only on days 14 and 21 after MCAO and on days 7 and 14 after PT. On close inspection, traditional pellet counts tended to overestimate motor abilities in mice before stroke, in case when they required a high number of reaches to retrieve individual pellets. For MCAO, we observed a progressive increase in reach duration accompanied by a decrease in hand movement speed. Deficits after PT were exemplified by the presence of hand slips throughout days 7 to 21. The high number of reaching attempts throughout the experiments confirmed that animals maintain their abilities to perform reaching movements with the arm and preserve high levels of task engagement. In summary, our results establish refined parameters for the quantification of differential post-stroke motor deficits in the two applied stroke models (Movie S1).

### Correlations of motor deficits with striatal or cortical lesion locations

We next addressed the contribution of lesion locations to the generation of motor deficits. Since both stroke groups showed comparable overall lesion volumes (Fig 2C), we pooled the data from both stroke groups for lesion-symptom correlation. Surprisingly, traditional pellet count showed no significant correlations with either total lesion volume or striatal or cortical ischemia (Fig. 4A). Instead, several other refined outcome parameters revealed lesion-specific correlations (Figs. 4B-D and S5). For example, the success coefficient on day 7 after stroke was significantly correlated with total lesion volume and cortical ischemia (p<0.05), but not with striatal ischemia (Fig. 4B). Slip depth highly correlated with cortical ischemia (p < 0.01) but did not reach the significance level for total or striatal lesion volume (Fig. 4C). The duration of reaching on day 21 was inversely correlated with striatal and cortical lesion volume (p<0.001 and p<0.01, Fig. 4D), showing slow movement after striatal and fast movement after cortical ischemia. In summary, lesion-symptom correlations provided additional evidence for brain-region-specific deficits following cortical or striatal ischemia in mice.

**Figure 4.**
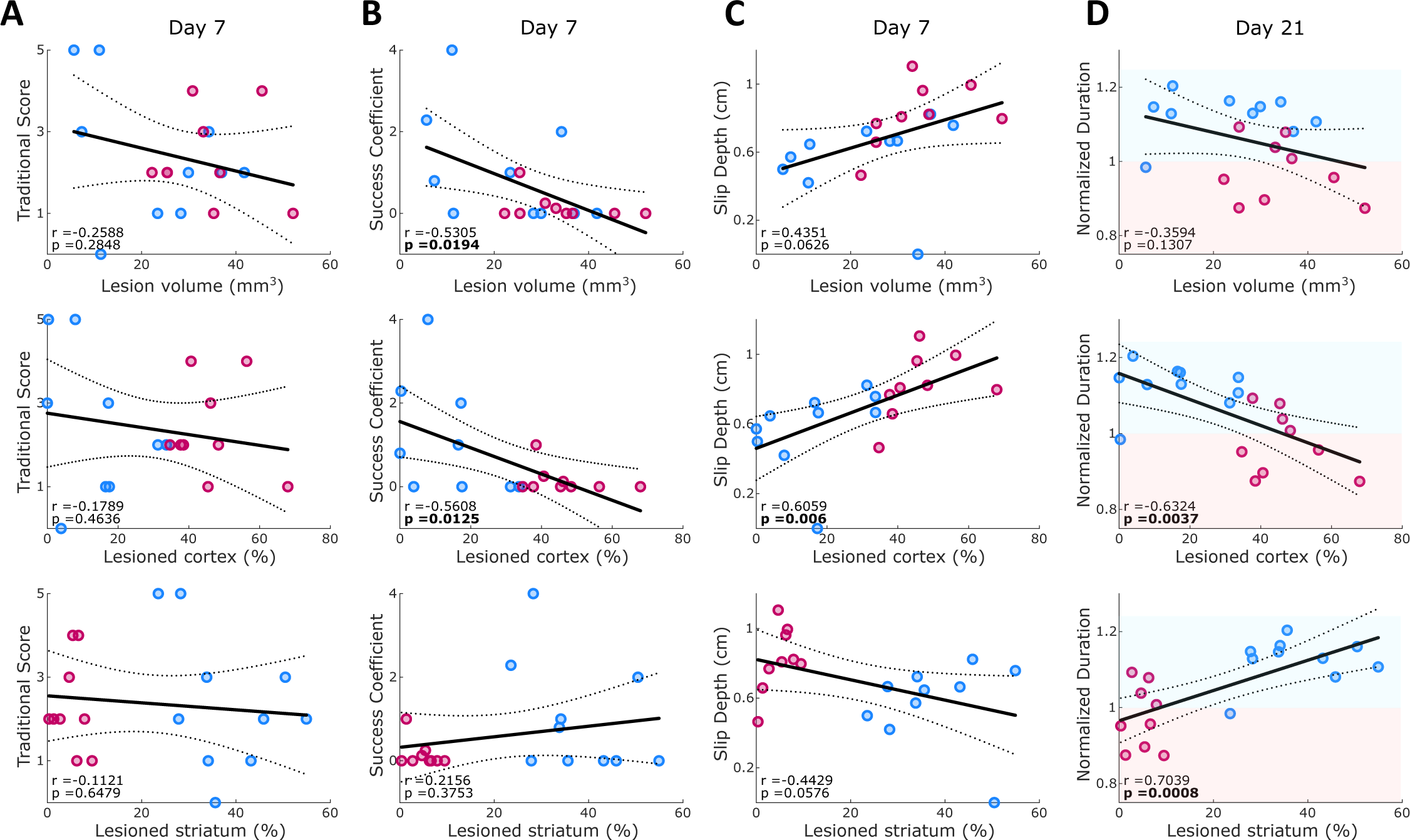
Correlation between outcome parameters and either total lesion volume, cortical or striatal lesion percentage. Correlations are reported for pooled dataset of MCAO and PT animals on day 7 after stroke for traditional pellet count (A), success coefficient (B) and slip depth (C), as well as on day 21 for normalized duration of the reaching cycle (D). Lines indicate linear fit and 95% confidence intervals. Red and blue shaded areas in D indicate decreased or increased reaching duration in comparison to pre-stroke behavior. Values report Pearson coefficients (r) and corresponding p-values. Significant correlations are marked in bold font for p<0.05.

### Compensatory changes in the ipsi-lesional hand after stroke

We finally utilized the bilateral video information to screen for kinematic changes of ‘non-affected’ ipsi-lesional hand in both stroke groups. Using LDA, we discovered changes after PT and MCAO, primarily in speed and distance-related kinematic parameters of reaching attempts (Figs. 5A-C). LD1 explained 59.9% of the variance in the dataset, followed by LD2 with 15.2%. Changes in LD1 and LD2 were significant for both stroke groups (p < 0.001), with significant differences in LD1 scores for days 14 and 21 in PT, and day 21 in MCAO animals (Fig. 5B). Analysis of individual parameters showed significant changes in the PT group for duration, hand acceleration, and path length (Fig. 5D). These movement adaptations after PT occurred when animals dropped in reaching performance not only in the contra-lesional, but also in the ipsi-lesional hand on day 7 after stroke (Fig. S2B). Different form the contra-lesional hand, the ipsi-lesional hand regained pre-lesion levels of reaching success from post-stroke day 14 onwards. Our observations, therefore, highlight potential compensatory movement strategies of the ipsi-lesional hand for reestablishing unilateral reaching success after stroke in mice.

**Figure 5.**
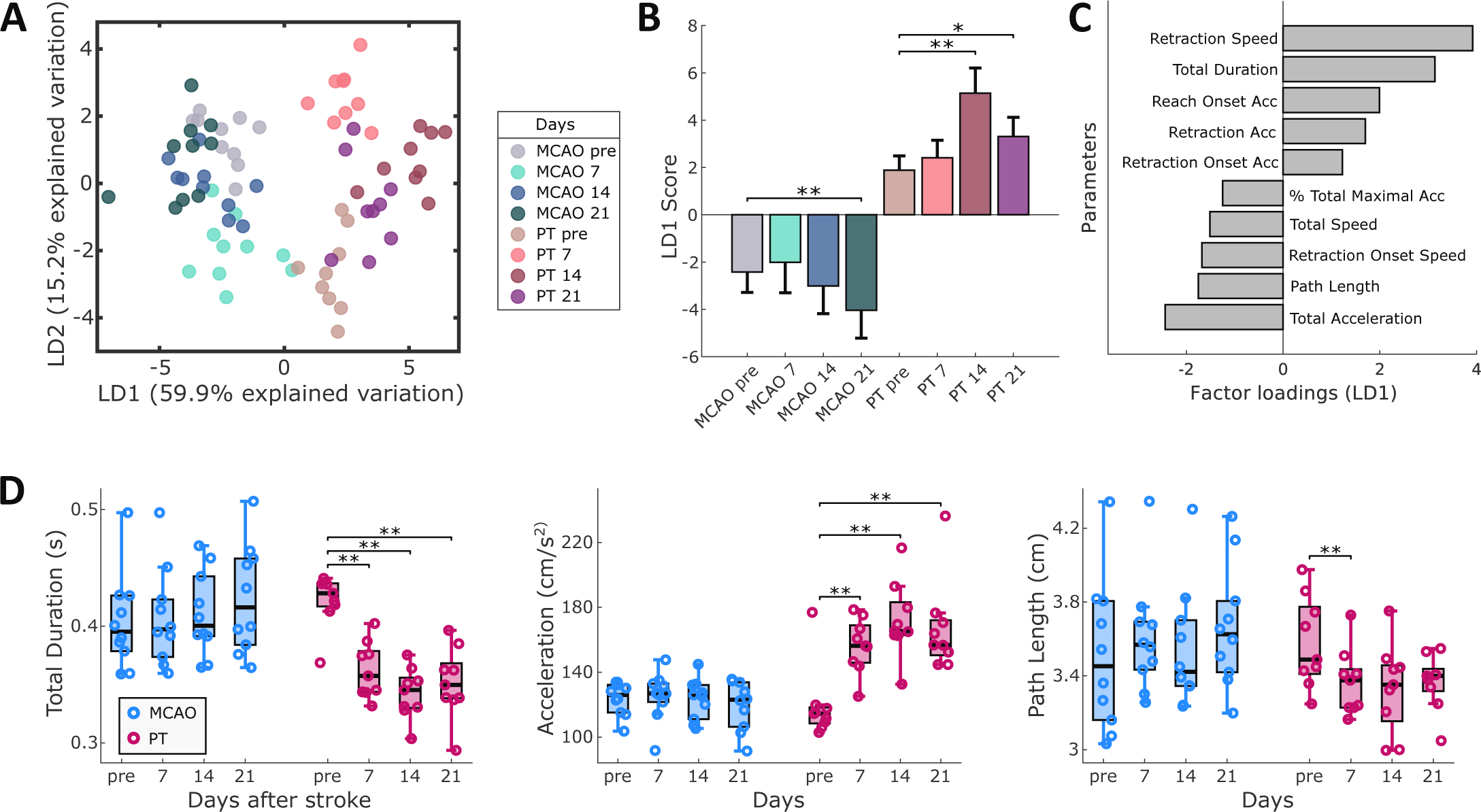
Compensatory kinematic changes in the ipsi-lesional hand. (A) Linear discriminant analysis (LDA) identifies the presence of ipsi-lesional motor adaptations, following MCAO and PT. (B) LD1 scores show significant changes on different recording days for both stroke groups. (C) Speed and distance-related parameters primarily account for the observed movement changes, as shown by the top 10 factor loadings on LD1. (D) Individual parameters provide information on new movement strategies after PT, composed of faster and shorter reaching movements. Bar graphs are reported as mean ± SD and box plots with Tukey method, *p<0.05, **p < 0.01.

## Discussion

In the present work, we developed a toolbox for the refined analysis of motor deficits in the Staircase test. In a proof of concept, we show that this toolbox can detect differential motor deficits following either MCAO or PT in mice. Additionally, lesion location in the striatum was linked to a progressive symptom manifestation of bradykinesia. In contrast, ischemia in cortex was selectively correlated with hand slips from the stabilizing staircase platform. In both stroke models, the number of retrieved pellets was altered, but this traditional measure did not correlate with stroke volume or the amount of cortical or striatal ischemia. Instead, we were able to describe selective motor features that showed significant correlations with lesion location. Together, these results provide evidence for the benefit of detailed behavioral profiling for translational stroke research.

### Lesion-symptom correlations

Several studies in rodents have shown correlations between lesion volume and global outcome parameters. In these studies, the effect of moderately sized lesions was compared to hemispheric infarcts covering most of the striatal and cortical brain tissue.^5,12^ However, large hemispheric strokes are suboptimal for animal research due to sickness behavior, which may interfere with biological recovery and the evaluation of sensorimotor function.^18^ Under ethical considerations, reducing lesion sizes in mice is of interest for 3R principles. Additionally, most strokes that cause permanent deficits in humans are in the range of small to medium-sized infarcts.^36,37^ For moderately sized infarcts in rats, traditional pellet count was not a fine-grained enough measure to distinguish between cortical or striatal ischemia.^12^ These findings agree with our present results in mice. Instead, refined movement analysis augmented the sensitivity of symptom correlations with small differences in infarct volume.

### Basal ganglia and stroke deficits

In humans, the location of ischemia in either the cortex or basal ganglia leads to differences in motor deficits, outcome prediction, and treatment response.^7–9,11,38,39^ Miyai et al. proposed that the reduced recovery of basal ganglia stroke compared to isolated cortical ischemia is due to distorted communication in corticothalamic-basal networks for motor learning.^38^ Physiologically, this is plausible, as identified structures, such as the dorsolateral striatum, are implicated in motor learning^40^, and stroke recovery in mice is accompanied by the reformation of coordinated neural activity in cortical and striatal ensembles.^41^ Our results show that striatal stroke in mice causes distinct behavioral deficits with an altered time course of manifestation. These insights support the concept that stroke in the basal ganglia deserves to be recognized as a distinct entity for the purposes of recovery and post-stroke therapy.

### Cortex and stroke deficits

A human stroke that affects structures along the corticospinal tract leads to severe hemiparesis.^9,10^ In agreement with previous literature^44^, our results showed that post-stroke deficits in mice after cortical ischemia did not mirror the severity of clinical symptoms in arm movements. These differences have been attributed to the divergence of the functional roles of the corticospinal tract in rodents, non-human primates, and humans.^45,46^ Independently of this discussion, our results establish motor profiles that are unique to mouse models of cortical stroke.

### Motor deficits vs. compensatory movements

We interpret the slow increase in reach duration after MCAO, over a period of three weeks, as a progressive motor symptom rather than a compensatory movement. Compensation is present when new motor patterns emerge that improve post-stroke performance, but the original movement does not recover.^16,42,43^ In our MCAO experiments, a decrease in reach duration developed along with declining performance in several outcome measures, including traditional pellet count and success coefficient. Therefore, our analysis provides a refined view on the distinction between prolonged symptom manifestation and compensation. This capability was further exemplified by progressive changes in movement patterns of the paw on the ‘non-affected’, ipsi-lesional hand after cortical photothrombosis (PT).

Over several weeks, mice after PT progressively reached faster with the ipsi-lesional hand and were able to fully recover their ipsi-lesional reaching success to pre-stroke levels. We interpret the faster reaching on the ipsi-lesional hand as a possible compensation for the prominent deficits of the contra-lesional hand, where hand slips indicate an inability to establish a firm grip onto the stabilizing staircase platform. These results also raise the notion that Staircase performance requires bilateral coordination. The functional evaluation of either side of the body cannot fully be discerned in the test. In experimental cases where the isolation of laterality plays a more central role, the single pellet reaching task may present a resort strategy.

### Limitations of the study

While our study presents a comprehensive analysis of forelimb movements in ischemic stroke models, there are several potential limitations that merit discussion. The Staircase task design inherently introduces variability based on factors such as the positioning of the body with respect to the hand, which we did not analyze in the present study. Body position may influence kinematic parameters and overall motor coordination. Future studies might benefit from separate analyses, also accounting for body position variations or shoulder and elbow tracking. Moreover, vertical hand slips could occur as part of the locomotion into the Staircase box or as part of the bilateral hand coordination to achieve the reaching attempts. Therefore, hand slips could further be delineated into different categories in future iterations of refined motor analysis. As a second point, the application of dimensionality reduction techniques such as LDA requires careful consideration due to its potential to introduce variability based on noise in the dataset, rather than meaningful deficits. However, in our dataset, the post-hoc analysis of the most correlated motor features in LDA revealed several significant and meaningful motor deficits. We therefore reason, that LDA served as an appropriate analysis tool for post-hoc analysis. Lastly, we performed correlations between lesion locations and outcome parameters in combined datasets of the two stroke models. We judge this approach as justifiable because both stroke models showed similar total lesion volumes (Fig 2C). Also, we did not observe any indication of a Simpson paradox, as correlation trends were maintained when looking at each stroke group, separately. To support this second argument, we also kept distinct colors for the MCAO and PT groups for the correlation plots in Fig. 4 and S5 to allow for easier visual separation.

### Conclusion

In summary, our results show that the use of machine learning for tracking hand and pellet motions in the Staircase can reveal differential motor deficits in commonly used mouse models of stroke. The algorithms from this study are freely available for download as a toolbox, *MouseReach*, from online repositories (see methods). These developments may foster improved validity of translational research. Parallel clinical research efforts are underway to establish wearable technologies for improved monitoring of motor scores, such as the Fugl-Meyer assessments in stroke patients.^47,48^ These combined developments may serve as important step towards symptom-specific understanding and personalized treatment in stroke research.

## Acknowledgements

We would like to acknowledge the excellent technical help provided by R. Gomaid, Z.E. Braun, and M. Renz for data collection and care of the mice. We thank A. Schill, A. von Garnier and J. Ode from the Scientific Laboratories Center at Charité for their technical work in customizing the Staircase hardware and the Service Unit Biometry at Charité for their advice on statistical planning.

## Author contribution statement

M.S. and N.W. created MouseReach. M.S., R.D.S., J.W., J.L., and L.M. conducted experiments. M.S. and R.D.S. analyzed data. S.M. and P.B.S. aided in MRI lesion analysis. R.P. provided technical support. R.S. and S.D. contributed to coding and project initiation. C.H., M.E., U.D., and A.V. participated in discussions. N.W. and M.S. authored the manuscript. N.W. and C.H. conceived the study. All authors have read and approved the final version.

