## Supplementary Figures for "Refined movement analysis in the Staircase test reveals differential motor deficits in mouse models of stroke"

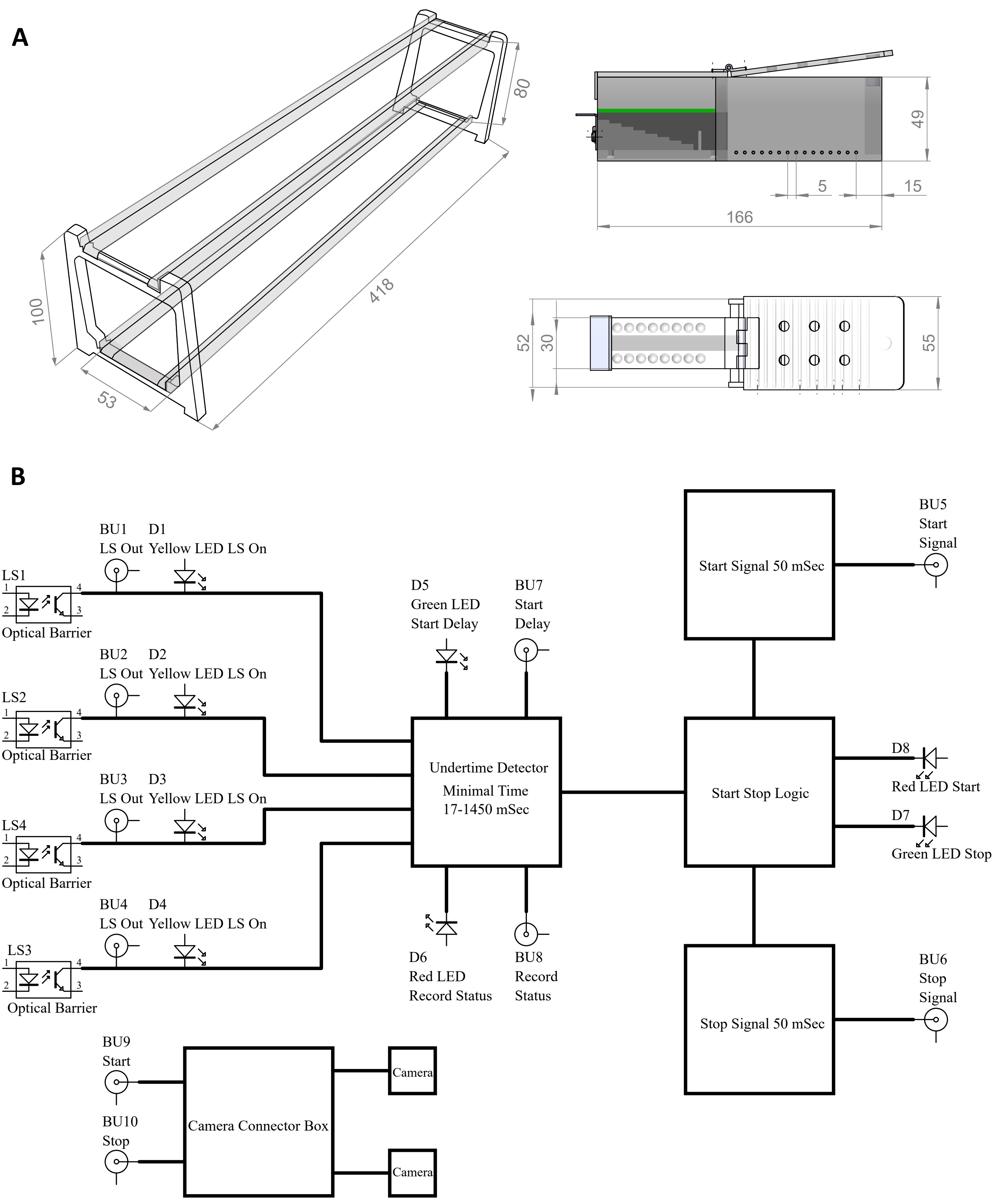


**Supplementary Figure 1. Hardware used in the multi-Staircase setup.** (A) Schematics of the positioning platform used for placement of four Staircases boxes stacked vertically onto two levels and a single Staircase box with side and top views. All values are displayed in millimeters. (B) Schematic of trigger hardware with optical barriers that enables instantaneous activation of both cameras upon mouse entry into the reaching chamber as well as deactivation after exiting the chamber.

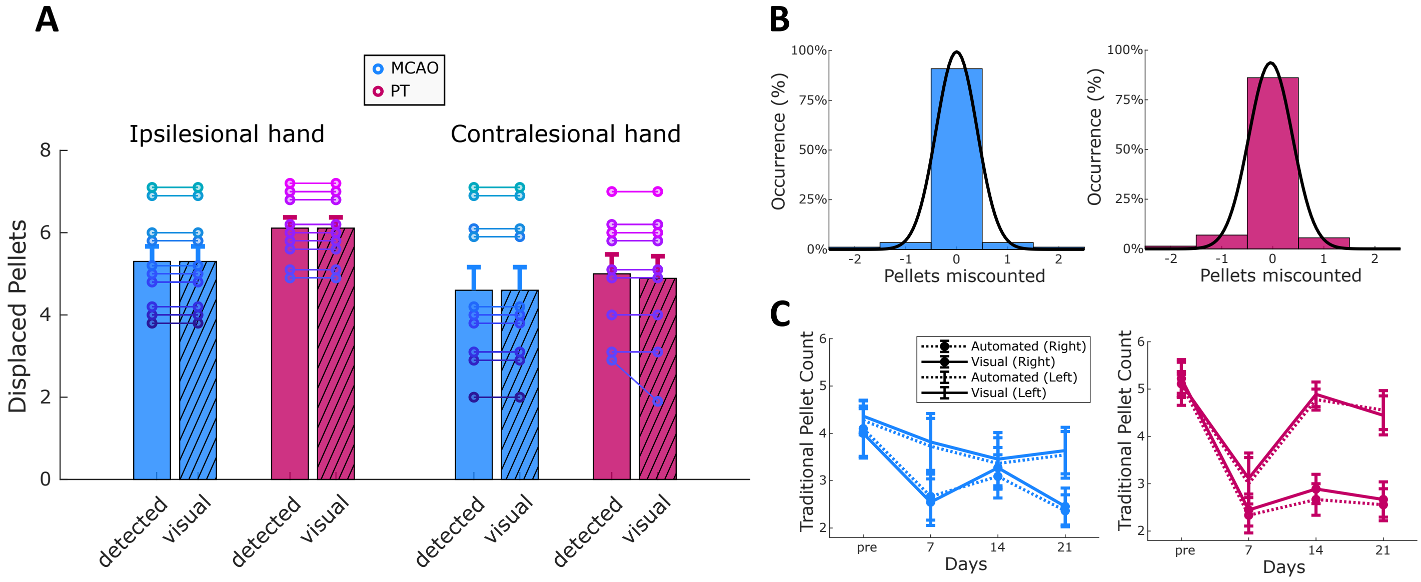


**Supplementary Figure S2. Validation of performance for detection of pellet removals and pellet count.** (A) Automatically detected pellet removals were compared to visual annotation by blinded raters for MCAO (blue, n=10 mice) and PT groups (magenta, n=9 mice). Individual values per animal and group means are shown for a single experimental day. Once a pellet was removed, there existed the possibility that it was either eaten by the mouse or dropped out of the hand and landed back onto another stair, thereby contributing to the final pellet count. (B) Gaussian curve that illustrates the marginal error rate for automated detection of pellet removals. Data are pooled for all MCAO and PT animals across all recording days (pre-stroke and day 7, 14, 21 after stroke). (C) Comparison of the traditional pellet count computed with MouseReach and via visual annotations.

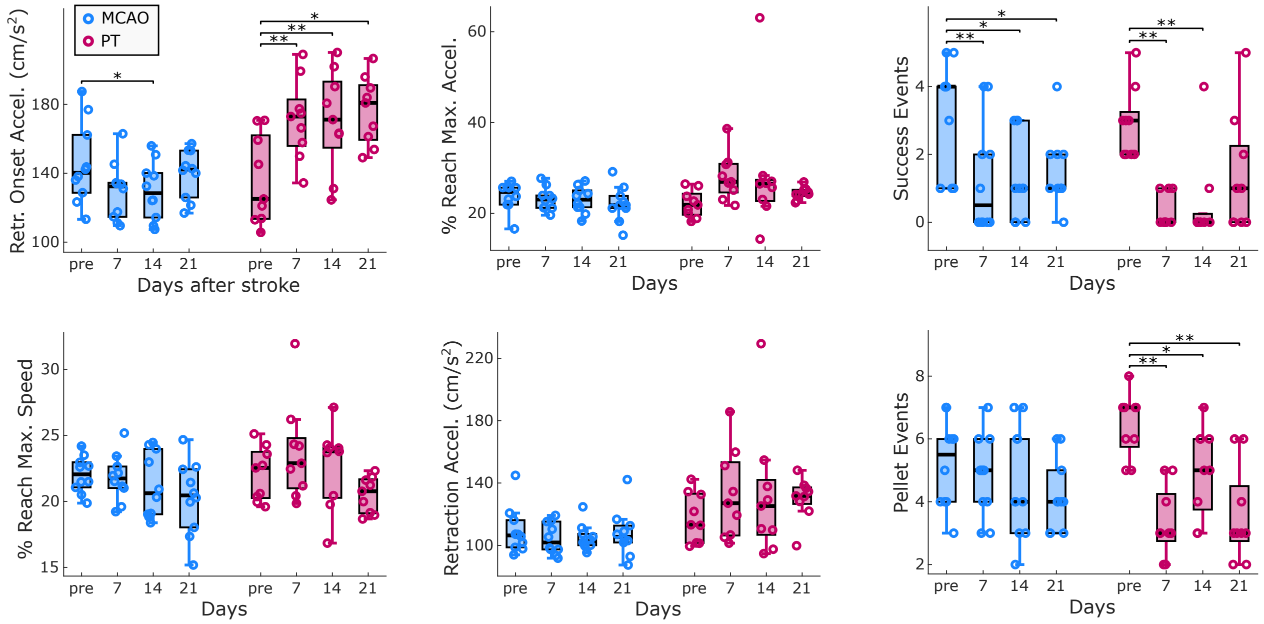


**Supplementary Figure S3.** Statistical analysis of outcome parameters reported among the top ten LD1 factor loadings, that have not been reported in main Fig. 3C due to space constraints. Pellet events correspond to the amount of sugar pellets displaced from the staircase, that were then either eaten or dropped. Success events count the total number of pellets that have been eaten after pellet displacement. Box plots are reported with Tukey method, *p<0.05, **p < 0.01.

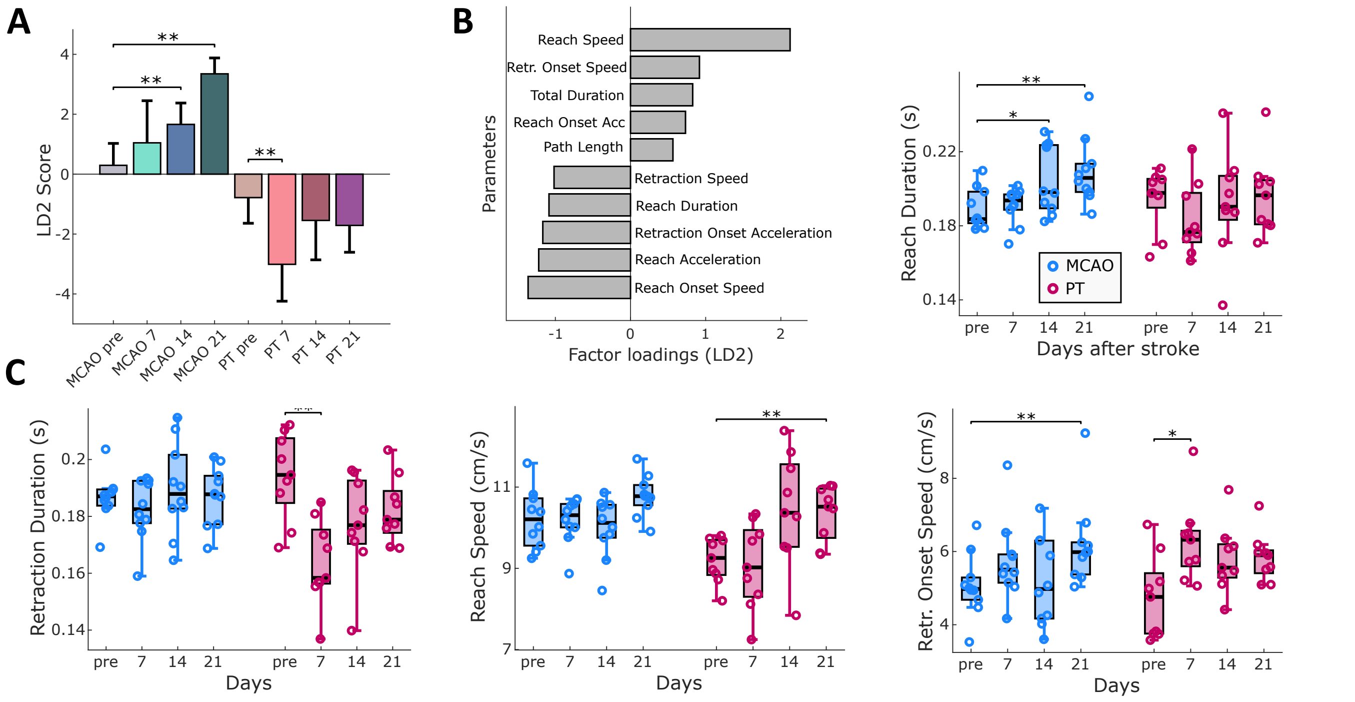


**Supplementary Figure S4.** Results for linear discriminant 2 (LD2), for post-stroke changes in the contralesional hand (complementing results from main Fig. 3 on LD1). (A, B and C) show LD2 scores, top ten factor loadings, and the results for individual selected outcome parameters. LD2 factor loadings captures speed-related changes, but not global parameter changes. Like LD1, changes in LD2 scores evolved with a delay after MCAO or presented immediately after PT. Reach duration captures the duration that the hand requires from the timepoint of movement initiation to the maximum extension of the hand towards the pellet. Bar graphs are reported as mean ± SD and box plots with Tukey method, $*p<0.05$, $**p < 0.01$.

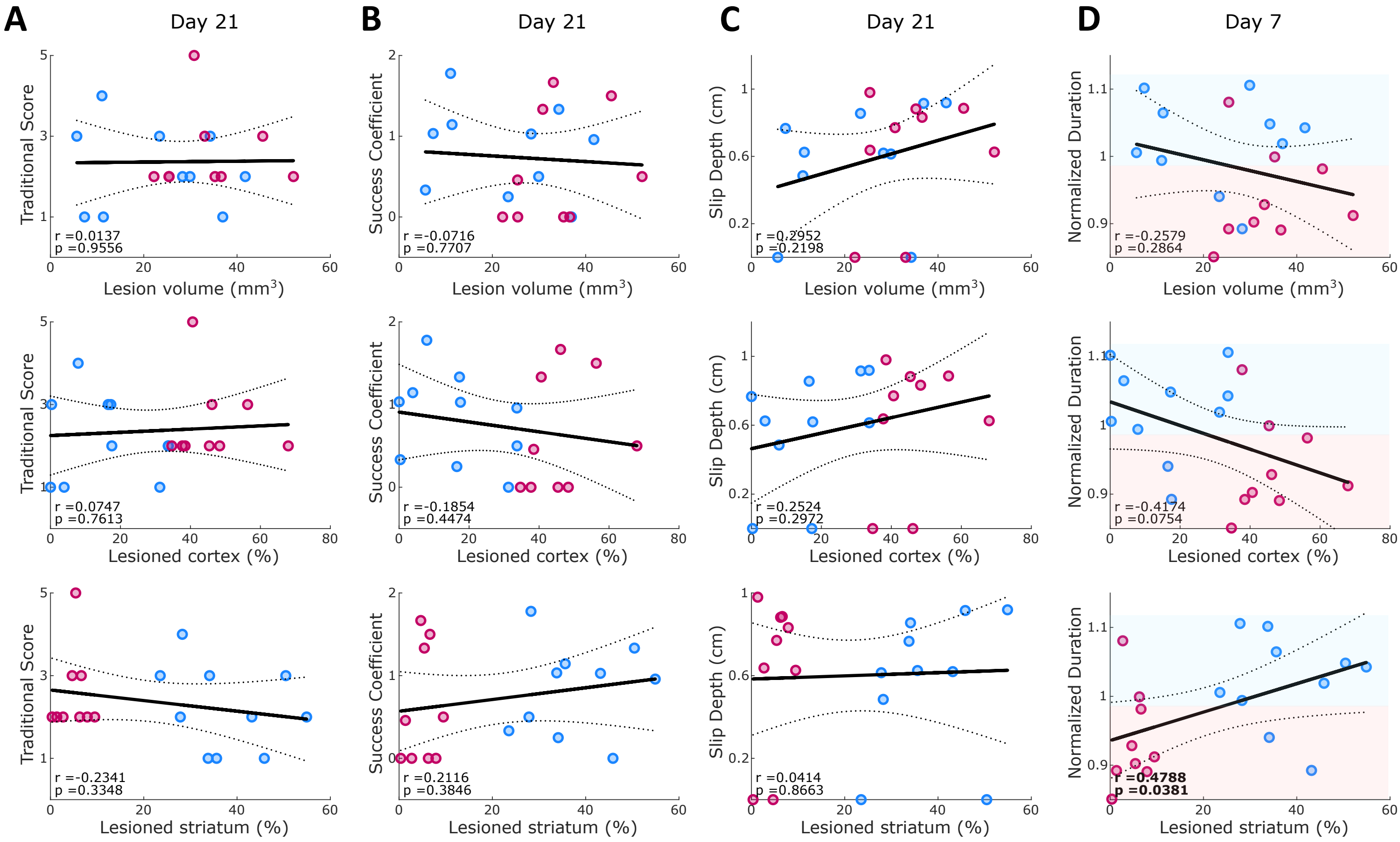


**Supplementary Figure S5.** Complementary timepoints for lesion-symptom correlations that are shown in main Fig. 4. Data are shown for post-stroke day 21 in MCAO and PT mice for traditional pellet count (A), success coefficient (B), slip depth (C), as well as the duration of the entire reach cycle on day 7 (D). Lines indicate linear fit and 95% confidence intervals. Red and blue shaded areas in D indicate decreased or increased reaching duration in comparison to pre-stroke behavior. Values reported include Pearson coefficients (r) and corresponding p-values. Significant correlations are marked in bold font for $p<0.05$.

**Supplementary Table S1**: List of 30 parameters for quantification of task performance

**Kinematic features**

| **Time (s)** | **Acceleration (cm/s$^2$)** |
|---|---|
| Total Duration | Total Acc |
| Reach Duration | Reach Acc |
| Retraction Duration | Retraction Acc |
| Slip Time | Reach Onset Acc |
| Grab Time | Retraction Onset Acc |
| **Position [cm]** | **Cycle Fraction (%)** |
| Path Length | Max Speed |
| Reach Distance | Reach Max Speed |
| Slip Depth | Retraction Max Speed |
| Grab Depth | Max Acc |
| **Speed (cm/s)** | Reach Max Acc |
| Total Speed | Retraction Max Acc |
| Reach Speed | |
| Retraction Speed | |
| Reach Onset Speed | |
| Retraction Onset Speed | |

| **Event related Information** |
|---|
| Success Coefficient |
| Success Events |
| Reach Events |
| Pellet Events |
| Slip Events |

Abbreviations: Acc (acceleration), Dec (deceleration), Max (maximum). Kinematic features denote quantification of reach cycles, except for two parameters: ‘Slip Time’ and ‘Slip Depth’. The total reach cycle is further divided into a reach-phase towards the pellet, and a retraction-phase back to the starting position. The category ‘Cycle Fraction’ denotes the timepoint when each feature occurred within a reaching cycle, normalized to 100% from cycle onset to end.
